# Differential action modes of Neutrophil Extracellular Trap-targeted drugs define T cell responses in SARS-CoV-2 infection

**DOI:** 10.1101/2024.06.30.601403

**Authors:** Caio Santos Bonilha, Flavio Protasio Veras, Anderson dos Santos Ramos, Giovanni Freitas Gomes, Robertha Mariana Rodrigues Lemes, Eurico Arruda, José Carlos Alves-Filho, Thiago Mattar Cunha, Fernando Queiroz Cunha

**Affiliations:** Center for Research in Inflammatory Diseases, Ribeirão Preto Medical School, University of Sao Paulo, 14049-900, Brazil; Institute of Infection, Immunity and Inflammation, University of Glasgow, G12 8TA, UK; Institute of Biomedical Sciences, Federal University of Alfenas, 37130-001, Brazil; Virology Research Center, Ribeirão Preto Medical School, University of São Paulo 14049-900, Brazil

## Abstract

Neutrophil extracellular traps (NETs) play a dual role in SARS-CoV-2 infection, aiding early immune defense but also contributing to lung damage. While NET targeting may improve clinical outcomes in SARS-CoV-2 infection, its impact on adaptive immunity, crucial for fighting the virus, remains unclear. Our study demonstrates that both recombinant human DNase (rhDNase), degrading NET structure, and GSK484, inhibiting NET formation, reduce lung NET concentration and improve clinical outcomes in infected mice, yet they differ in their influence on T cell responses. We show that rhDNase does not impact T cell responses, whereas GSK484 diminishes virus-specific T cell responses. *In vitro*, GSK484 decreases dendritic cell antigen presentation by impairing antigen uptake and reduces IL-2 signaling by affecting its production by T cells. In a model of lung inflammation, GSK484 diminishes antigen-specific T cell activation and proliferation, while rhDNase shows a potential to boost T cell responses via the presence of NET fragments that reduce T cell activation threshold. Our findings suggest that NET targeting with rhDNase or GSK484 holds therapeutic potential for treating SARS-CoV-2 infection, while their distinct modes of action shape T cell responses during the infection.

## Introduction

The current era of coronavirus disease 2019 (COVID-19), marked by widespread population immunity resulting from prior infection and/or vaccination, has witnessed a substantial reduction in overall disease severity since the pandemic’s onset [1]. However, effective treatments remain crucial in managing the disease, particularly for vulnerable populations, including those who are unvaccinated, incompletely vaccinated, or vaccinated but immunocompromised [2].

The significance of adaptive immune responses in COVID-19 is becoming increasingly evident [2,3]. Within the spectrum of adaptive immune responses, the role of T cell responses emerges as vital in fighting the virus. Evidence suggests that individuals with robust, virus-specific CD8 [4] and CD4 [5] T cell responses experience faster clearance of severe acute respiratory syndrome coronavirus 2 (SARS-CoV-2) and better COVID-19 outcomes. In addition, SARS-CoV-2-specific T cells responses have been positively correlated with memory B cell responses and SARS-CoV-2-neutralizing antibody production [6]. Accordingly, therapeutic strategies should not only aim at alleviating clinical symptoms but also consider their repercussion on adaptive immunity [3].

In addition to cellular responses, emerging evidence underscores the role of neutrophil extracellular traps (NETs) in COVID-19 pathogenesis [7]. While NETs, composed of DNA and neutrophil granule proteins, serve as a physical barrier against SARS-CoV-2 spread [8], they also contribute to lung tissue damage by inducing airway epithelial cell apoptosis [9]. This deleterious effect has prompted interest in investigating NETs as potential targets for treating SARS-CoV-2 infection. However, NET targeting might alter adaptive immune responses [7], as NET components can alert the immune system to the presence of the pathogen and augment its response [10–13]. As such, an optimal NET-targeted therapeutic approach should aim to mitigate the detrimental effects of NETs while preserving or enhancing virus-specific immunity.

While NET targeting may improve clinical outcomes led by the infection [14], its effects on adaptive immune responses against SARS-CoV-2 remain unclear. We analyzed the impact of recombinant human DNase (rhDNase), which degrades the DNA backbone of NETs, and GSK484, which inhibits protein arginine deiminase 4 (PAD4)-dependent NET formation (NETosis) in a COVID-19 model. Both drugs improved clinical outcomes; however, we show that GSK484 reduces the function of dendritic cells (DC), crucial for inducing adaptive immunity, as well as T cell responses, while rhDNase does not affect virus-specific T cell responses. Our findings suggest that rhDNase and GSK484 hold promise for treating SARS-CoV-2 infection, while their distinct modes of action shape virus-specific T cell responses.

## Methods

### Sex as a biological variable

Our study utilized both men and women as human donors and male and female animals. Similar findings were observed irrespective of sex.

### Mice

K18-hACE2, OT-II, CD45.1 and C57BL/6 mice were bred in-house (Animal Breeding Center, Ribeirao Preto Medical School). All mice were of the C57BL/6 genetic background and specific pathogen free, maintained in accordance with local and federal regulations.

### COVID-19 model

SARS-CoV-2 virus (SPBR-02/2020 strain) that was titrated by plaque-forming units (PFU) assays using VERO E6 cells [15] was used to infect K18-hACE2 animals. Mice were intranasally instilled with 10^3^ PFU SARS-CoV-2 and maintained in a Biosafety Level 3 (BSL-3) laboratory. Body weight was assessed at baseline and on all days post-infection (dpi). Clinical scores were determined as previously described [14]. Whole lungs, mediastinal lymph nodes (mLN), and spleens were digested with 1 mg/mL Collagenase D (Sigma-Aldrich, #11088866001) [16], before flow cytometry analysis. For the analysis of SARS-CoV-2-specific T cells, lymphoid organ cells were pulsed with a SARS-CoV-2 peptide pool (Miltenyi Biotec, # 130-127-309) (1 μg/mL) for 30 minutes and then incubated with a Protein Transport Inhibitor (Thermo Fisher, #00-4980-03) for 6 hours.

### Lung inflammation model

Lymph node (LN) and spleen cells from OT-II mice were labelled with CFSE (Thermo Fisher, #C34554) (1.25 μM) and adoptively transferred (intravenously) into CD45.1 recipients, as previously described [17]. One day later, recipient mice were intranasally challenged with 10 μg Lipopolysaccharides (LPS) (Sigma-Aldrich, #L4516) and 10 μg ovalbumin (OVA) (Sigma-Aldrich, #A5253). Three days after challenge, mLNs were harvested, and processed for flow cytometry analysis.

### Treatments

Mice were treated subcutaneously with rhDNase (Roche, Pulmozyme) (10 mg/kg), or intraperitonially with GSK484 (MedChem Express, #HY-100514) (4 mg/kg), anti-TNF-α (Bio X Cell, #BE0058) (100 μg/mouse), or an isotype control (Bio X Cell, #BE0088) (100 μg/mouse). The first doses were given 30 minutes before intranasal instillations.

### DC culture

Bone marrow derived-DCs (BMDCs) were prepared using 10 ng/mL recombinant GM-CSF (Biolegend, #576306), as previously described [18]. On day 8, CD11c+ cells were isolated (Miltenyi Biotec, #130-125-835) and stimulated with 100 ng/mL TNF-α (Biolegend, #575206). Day-9 BMDCs were used for functional assays. Alternatively, splenic DCs were used, after isolation with a Miltenyi Biotec kit (#130-100-875). For antigen uptake assays, DCs were pulsed with Alexa Fluor 647 (AF647)-conjugated OVA (Thermo Fisher, #O34784) (20 μg/mL). DCs were pulsed with Eα_52-68_ (Anaspec, #AS-61621) or OVA_257-264_ (Anaspec, #AS-60193-1) peptide at 10 μg/mL, and then stained with the YAe (Thermo Fisher, # 11-5741-81) or 25-D1.16 (Thermo Fisher, #12-5743-81) antibody clones for the detection of MHC-II-or MHC-I-mediated antigen presentation, respectively. YAe and 25-D1.16 antibodies specifically bind to complexes of MHC molecules with their corresponding peptides, not to MHC molecules or free OVA peptides. Culture supernatants were used for detection of cytokines by ELISA.

### Mouse T cell culture

Isolated CD4 (Miltenyi Biotec, #130-104-454) or CD8 (Miltenyi Biotec, #130-104-075) T cells from C57BL/6 mice were stimulated with coated anti-CD3 (Thermo Fisher, #16-0032-85) (10 μg/mL) and soluble anti-CD28 (Thermo Fisher, #16-0281-86) (2 μg/mL). For the detection of intracellular cytokines, cells were re-stimulated with a Stimulation Cocktail (Themo Fisher, #00-4970-93) in the presence of a Protein Transport Inhibitor. Culture supernatants were used for detection of cytokines by ELISA.

### Human T cell culture

Peripheral blood mononuclear bells (PBMCs) were isolated from healthy donors using Histopaque-1077 (Sigma-Aldrich, #10771). Naïve T cells were then isolated (Miltenyi Biotec, #130-097-095), and from these cells, CD4 T cells were purified (Miltenyi Biotec, #130-094-131). The remaining fraction of naïve T cells was considered as naïve CD8 T cells. CD4 and CD8 T cells were labelled with CFSE and stimulated with coated anti-CD3 (Thermo Fisher, #16-0037-85) (10 μg/mL).

### DC-T cell co-culture

CFSE-labelled CD4 T cells from OT-II mice were cultured with BMDCs pulsed with OVA_323-339_ (Sigma-Aldrich, #O1641) (5 μg/mL), as previously described [19]. Culture supernatants were used for detection of cytokines by ELISA.

### NET isolation

Mouse neutrophils were isolated using Histopaque-1077 and Histopaque-1119 (Sigma-Aldrich, #11191). Cells were then stimulated with Phorbol 12-myristate 13-acetate (PMA) (Sigma-Aldrich, #P8139) (50 nM). After 3 hours, cells were harvested, centrifuged, and their supernatant was recovered. To increase NET concentration, supernatants were centrifuged at 18,000 xg for 10 minutes. For functional assays, NETs were used at 10 ng/mL, determined by the DNA concentration, measured by a NanoDrop Microvolume Spectrophotometer (Thermo Fisher).

### NET quantification

Samples were incubated in plates coated with anti-myeloperoxidase (MPO) (Thermo Fisher, #PA5-16672), washed, and DNA bound to MPO was quantified using a Quant-iT PicoGreen kit (Invitrogen, #P11496).

### Flow cytometry

Single-cell suspensions were incubated with 10% mouse serum to block Fc receptors before adding fluorochrome-conjugated antibodies, as previously described [20]. Viability was assessed using a fixable cell viability dye (eBioscience) according to the manufacturer’s instructions. For pSTAT5 detection, cells were fixed (BD Biosciences, #557870), and permeabilized (BD Biosciences, #558050), before being stained for flow cytometry analysis. For the detection of intracellular cytokines, transcription factors, or intrapeptidic citrulline (#ab100932), a Foxp3 Staining kit was used (eBioscience, #00-5523-00). Cells were run on a FACSCanto II or a FACSVerse (BD Biosciences) and analyzed using BCyto [21].

### ELISA

The concentration of cytokines and chemokines were determined using R&D Systems kits (DuoSet), according to the manufacturer’s instructions. SARS-CoV-2 N protein concentration was determined using a Proteintech kit (#KE30007).

### Statistics

Data is shown as mean ± SD. Statistical analyses were performed using Prism version 10 (GraphPad Software, Inc, CA, USA).

### Study approval

Experiments were performed in accordance with the guide for the use of laboratory animals of the University of Sao Paulo and approved by the ethics committee under protocol #066/2020. Written informed consent was obtained from all study participants prior to participation.

### Data availability

A Supporting Data Values file is available in accordance with JCI policy. Additional data and materials are available upon reasonable request.

## Results

### NET targeting with rhDNase and GSK484 improve clinical outcomes in SARS-CoV-2 infection

To assess the impact of NET-targeted drugs on adaptive immune responses during SARS-CoV-2 infection, we employed a murine model of COVID-19 (Figure 1A), which exhibits disease signs and lung pathology compatible with the human disease [22]. To target NETs, we used rhDNase, which inhibits NET-induced airway epithelial cell apoptosis by degrading NET DNA [9], and GSK484, a PAD4-selective inhibitor which has been shown to inhibit NETosis *in vitro* [23–27] and in models of lung inflammation [27–31]. Both rhDNase and GSK484 improved clinical disease (Figure 1B-C), showcasing the therapeutic potential of NET-targeted drugs for treating SARS-CoV-2 infection. We have previously shown that Cl-Amidine, a pan-PAD inhibitor prevents SARS-CoV-2-induced NETosis in human neutrophils [9]; however, hACE2 expression is limited to epithelial cells in the mice used in our model. Despite this, we found that both rhDNase and GSK484 decreased lung NET concentration (Figure 1D) without affecting neutrophil migration (Figure 1E-G), suggesting that NETosis in this model might be dependent on PAD4. Although no significant impact on the viral load was found (Figure 1H), we analyzed lung cytokines at 2 dpi to further understand the drugs’ effects on immune responses. Focus was given to cytokines secreted by cells such as DCs, the most specialized antigen-presenting cell (APC) that bridge innate and adaptive immunity. RhDNase, but not GSK484, significantly decreased the concentration of pro-inflammatory cytokines (IL-23, IL-1β, IL-6) at 2 dpi, while both treatments decreased IL-10 concentration and did not affect TNF-α concentration (Figure 1I). The effects of the drugs on cytokine secretion indicate potential impacts on immune responses.

**Figure 1.**
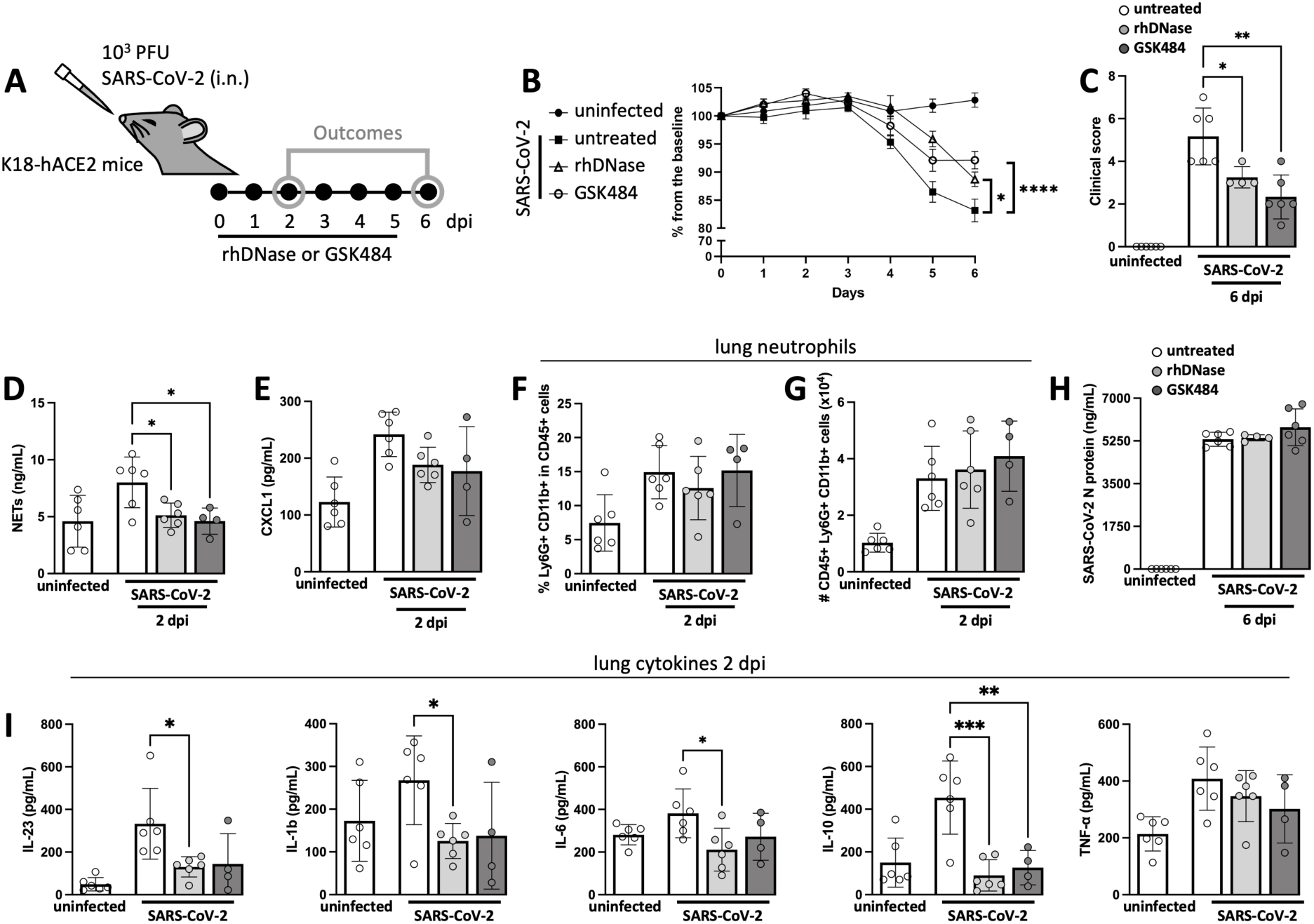
rhDNase I and GSK484 reduce clinical disease in a COVID-19 model. (**A**) K18-hACE2 mice (N=4-6) were intranasally infected with SARS-CoV-2 (10^3^ PFU) and treated daily with rhDNase or GSK484. (**B**) Body weight and (**C**) clinical scores were analyzed. (**D**) Concentration of NETs (MPO-DNA conjugates) and (**E**) CXCL1 in lung homogenates 2 days post-infection (dpi). (**F**) Percentage of Ly6G+ CD11b+ in CD45+ cells and (**G**) number of CD45+ Ly6G+ CD11b+ cells in the lungs 2 dpi. (**H**) Concentration of SARS-CoV-2 N protein in lung homogenates 2 and 6 dpi. (**I**) Concentration of cytokines in lung homogenates 2 dpi. Weight differences were compared by two-way ANOVA. All other statistical differences were determined by one-way ANOVA. ANOVA tests were followed by Tukey’s multiple comparison tests. *p<0.05, **p<0.01, ***p<0.001, and ****p<0.0001.

### GSK484 alters cDC responses during SARS-CoV-2 infection

Before evaluating the effects of the treatments on T cell responses, we assessed whether the drugs could influence cDC responses by analyzing lung and lymphoid organ cells at 2 dpi (Figure 2A). Neither drug affected the accumulation of APCs expressing low levels of CD11c in the lungs (Figure 2B-C). However, GSK484 decreased the accumulation of lung conventional DCs (cDCs) (Figure 2D-E), both type 2 (cDC2) (Figure 2F) and type 1 (cDC1) (Figure 2G), distinguished by their differential expression of CD103 [32]. Additionally, GSK484 decreased the expression of MHC-II in CD11c_high_ cells, suggesting a possible effect on lung cDC maturation (Figure 2H). Importantly, since the infection in this model does not induce the accumulation of conventional DCs (cDC) in the lungs—a possible sign of DC failure characteristic of COVID-19 [33–36]—the interpretation of the modulatory effects of the treatments requires caution. As such, these results suggest that GSK484 does not necessarily reduce cDC lung responses during the infection, but rather alters them.

**Figure 2.**
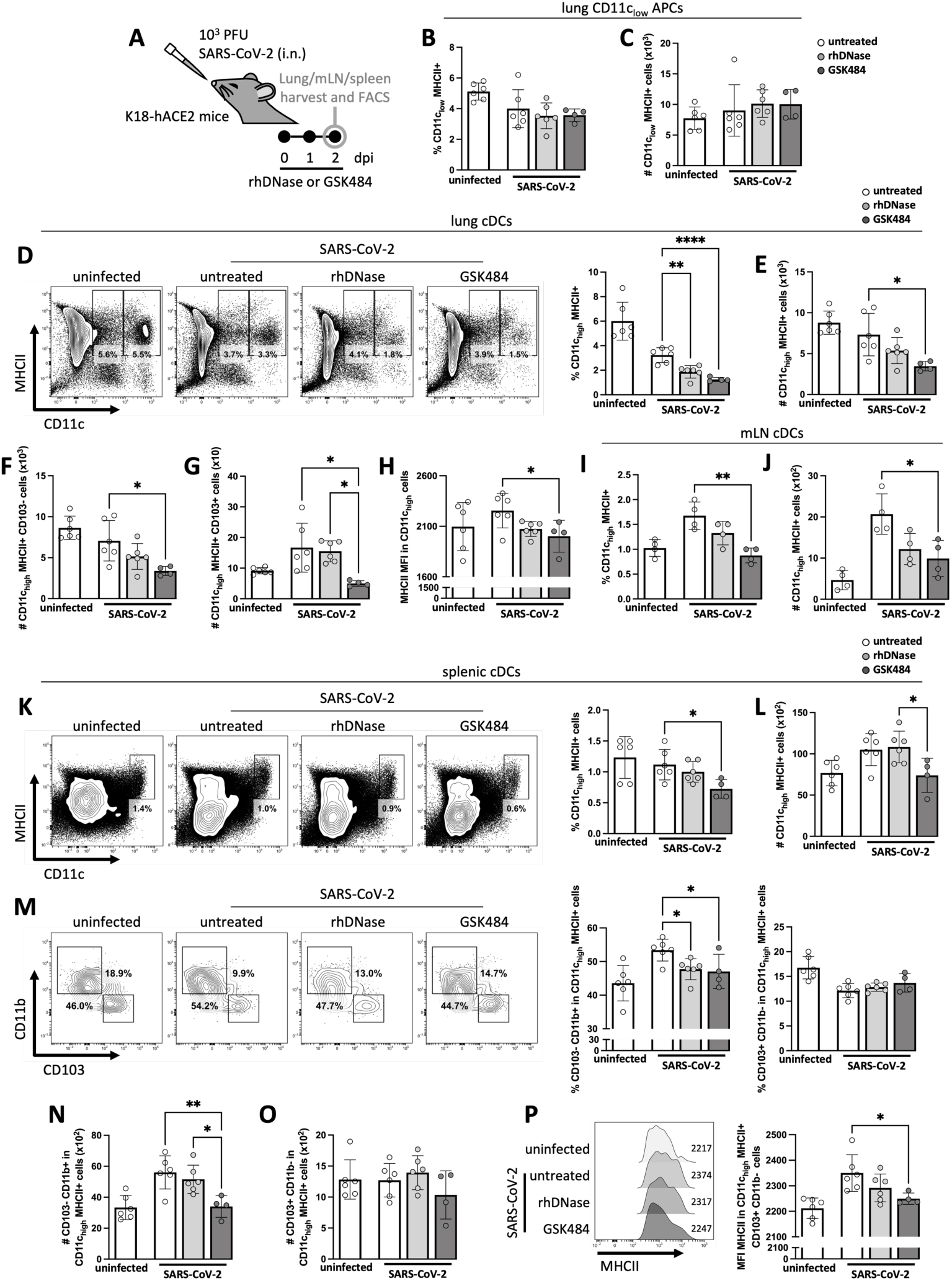
GSK484 affects cDC responses during SARS-CoV-2 infection. (**A**) Lungs, mediastinal lymph nodes (mLN), and spleens from SARS-CoV-2-infected mice (N=4-6) treated daily with rhDNase or GSK484 were harvested 2 days post-infection (dpi) for flow cytometry analysis. (**B**) Percentage and (**C**) number of CD11c_low_ MHC-II+ cells in the lungs. (**D**) Percentage and (**E**) number of CD11c_high_ MHC-II+ cells in the lungs. (**F**) Number of CD11c_high_ MHC-II+ CD103-and (**G**) CD103+ cells in the lungs. (**H**) MHC-II median fluorescence intensity (MFI) in CD11c_high_ cells in the lungs. (**I**) Percentage and (**J**) number of CD11c_high_ MHC-II+ cells in the mLNs. (**K**) Percentage and (**L**) number of CD11c_high_ MHC-II+ cells in the spleens. (**M**) Percentage of CD103-CD11b+ and CD103+ CD11b-in CD11c_high_ MHC-II+ cells in the spleens. (**N**) Number of CD11c_high_ MHC-II+ CD103-CD11b+ and (**O**) CD11b-cells in the spleens. (**P**) MHC-II MFI in CD11c_high_ MHC-II+ CD103+ CD11b-cells. Differences between infected groups were determined by one-way ANOVA, followed by Tukey’s multiple comparison test. *p<0.05, **p<0.01, and ****p<0.0001.

Both cDC1 and cDC2 can prime naïve T cells in the lungs before their migration to lymphoid organs [37]; however, classically, cDCs uptake antigen in affected tissues to prime T cells in lymphoid organs. Importantly, it has been demonstrated that in addition to the mLNs, the spleen can also serve as a primary site for the initiation of T cell responses to respiratory virus infections [38]. We show that GSK484 decreased the accumulation of cDCs in the mLNs (Figure 2I-J) and spleens (Figure 2K-L) induced by the infection. Within cDC subsets, we noticed that GSK484 decreased the accumulation of splenic cDC2 triggered by the infection (Figure 2M-N). While no effect was observed in the number of cDC1 in GSK484-treated mice (Figure 2O), these cells exhibited lower expression of MHC-II (Figure 2P). These findings indicate that GSK484 affects cDC responses during the infection, with potential implications to T cell responses.

### GSK484, but not rhDNase, diminishes SARS-CoV-2-specific T cell responses

Considering the impact of NET targeting with GSK484 on cDC responses during SARS-CoV-2 infection, we analyzed whether the drugs could affect local T cell responses. Lung T cell accumulation was analyzed at 6 dpi (Figure 3A), when a notable increase in activated T cell accumulation was evident, as compared with uninfected or mice infected at earlier time points (data not shown). GSK484 treatment led to a reduction in the percentage of activated T cells (Figure 3B), as well as the number of CD4 (Figure 3C) and CD8 (Figure 3D) T cells. The decline in T cell accumulation was not restricted to a particular subset, as cytotoxic (Figure 3E-F), Th17 (Figure 3I-J), and T_reg_ cells (Figure 3K-L) were affected. Additionally, we observed that GSK484, in contrast to rhDNase, significantly decreased the number of proliferating T cells in the mLNs (Figure 3M-N), indicating a potential effect on T cell proliferation.

**Figure 3.**
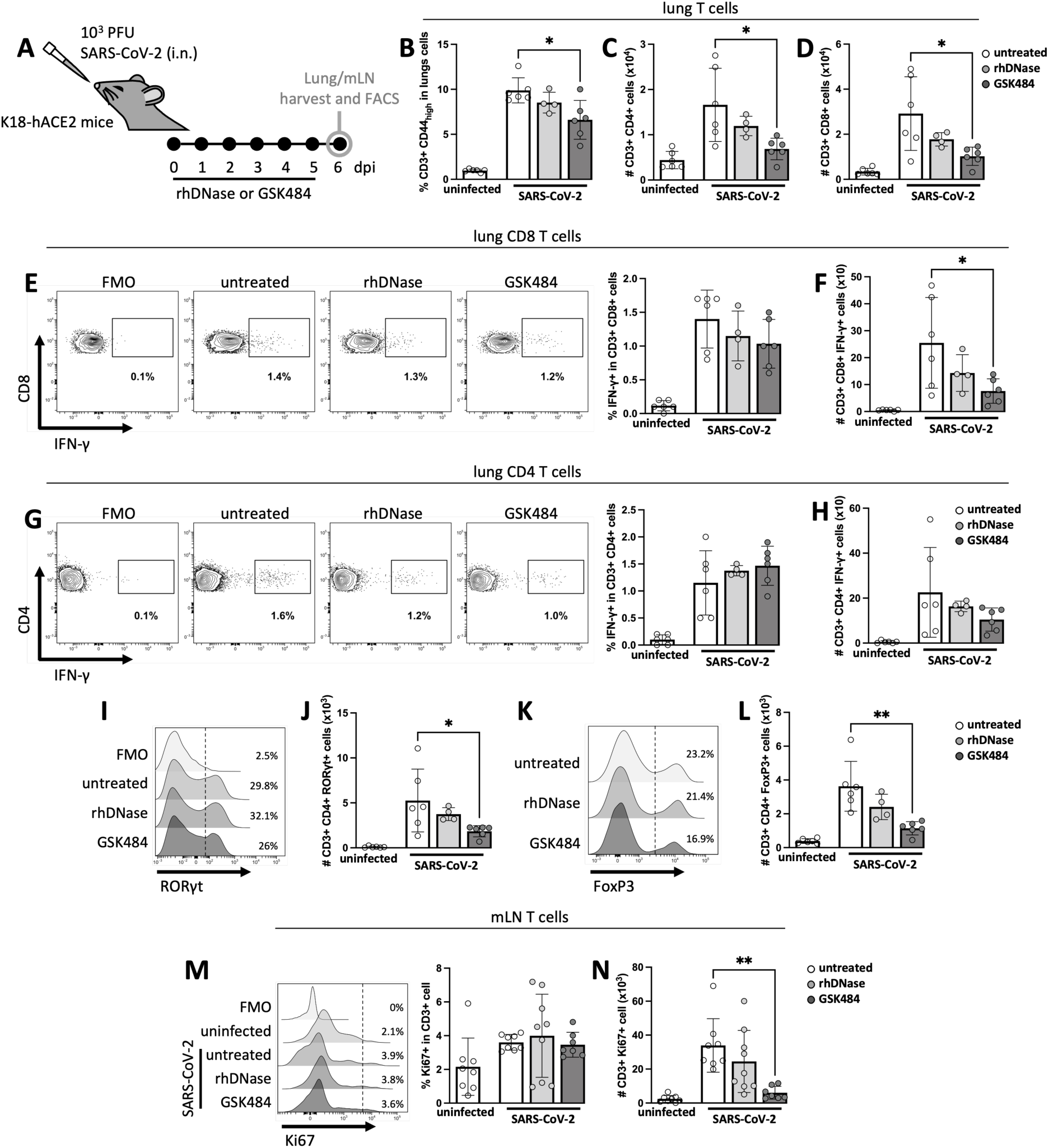
GSK484 limits lung T cell responses during SARS-CoV-2 infection. (**A**) Lungs and mediastinal lymph nodes (mLN) from SARS-CoV-2-infected mice (N=4-9) treated daily with rhDNase or GSK484 were harvested 6 days post-infection (dpi) for flow cytometry analysis. (**B**) Percentage of CD3+ CD44_high_ cells in the lungs. (**C**) Number of CD3+ CD4+ or (**D**) CD3+ CD8+ cells in the lungs. (**E**) Percentage of IFN-γ+ in CD3+ CD8+ cells and (**F**) number of CD3+ CD8+ IFN-γ+ cells in the lungs. (**G**) Percentage of IFN-γ+ in CD3+ CD4+ cells and (**H**) number of CD3+ CD4+ IFN-γ+ cells in the lungs. (**I**) Percentage of RORγt+ in CD3+ CD4+ cells and (**J**) number of CD3+ CD4+ RORγt + cells in the lungs. (**K**) Percentage of FoxP3+ in CD3+ CD4+ cells and (**L**) number of CD3+ CD4+ FoxP3+ cells in the lungs. (**M**) Percentage of Ki67+ in CD3+ cells and (**N**) number of CD3+ Ki67+ cells in the mLNs. Differences between infected groups were determined by one-way ANOVA, followed by Tukey’s multiple comparison test. *p<0.05, and **p<0.01.

Given the observed reduction in local T cell responses, we investigated the impact of NET targeting on virus-specific T cells. Cells harvested at 6 dpi from lymphoid organs containing APCs were re-stimulated *in vitro* with a SARS-CoV-2 peptide pool (Figure 4A). Virus-specific effector responses were evident when comparing cells from the infected group with and without re-stimulation. Notably, GSK484, but not rhDNase, significantly reduced the number of IFN-γ producer splenic CD8 (Figure 4B-D), splenic CD4 (Figure 4E-G), mLN CD8 (Figure 4H), and mLN CD4 (Figure 4I) T cells. These data suggest that unlike rhDNase, GSK484 diminishes SARS-CoV-2-specific T cell responses during the infection.

**Figure 4.**
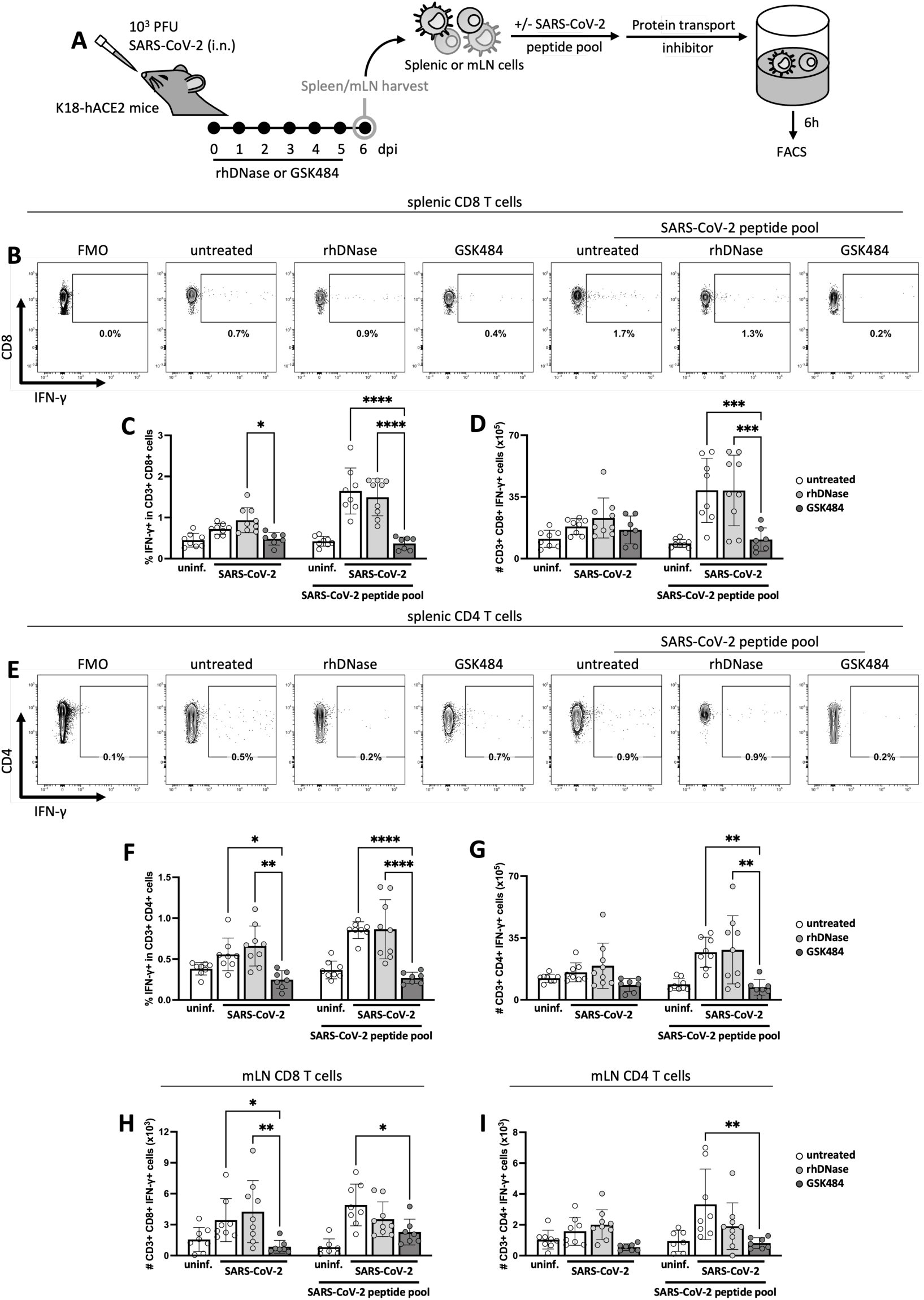
GSK484 diminishes SARS-CoV-2-specific T cell responses. (**A**) Spleens and mediastinal lymph nodes (mLN) from SARS-CoV-2-infected mice (N=7-9) treated daily with rhDNase or GSK484 were harvested 6 days post-infection (dpi). Cells were then pulsed with a SARS-CoV-2 peptide pool, treated with Brefeldin A, and analyzed 6 hours later by flow cytometry. (**B-C**) Percentage of IFN-γ+ in CD3+ CD8+ cells and (**D**) number of CD3+ CD8+ IFN-γ+ cells in the spleens. (**E-F**) Percentage of IFN-γ+ in CD3+ CD4+ cells and (**G**) number of CD3+ CD4+ IFN-γ+ cells in the spleens. (**H**) Number of CD3+ CD8+ IFN-γ+ and (**I**) CD3+ CD4+ IFN-γ+ cells in the mLNs. Differences between infected groups were determined by two-way ANOVA, followed by Tukey’s multiple comparison test. *p<0.05, **p<0.01, ***p<0.001, and ****p<0.0001.

### GSK484 minimally impacts DC maturation but impairs antigen presentation, resulting in reduced T cell priming

As our investigation revealed notable effects of GSK484 on both DC and T cell responses during SARS-CoV-2 infection, we examined its impact on these cells *in vitro*. GSK484 inhibits PAD4, an enzyme that catalyzes the post-translational modification of arginine to citrulline. This inhibition blocks NET formation by preventing histone citrullination, a key step in PAD4-dependent NETosis [39–41]. Nevertheless, citrullination can modulate the function of several molecules, which suggests that PAD4 inhibition might affect the function of other proteins involved in immune responses [42].

To examine the impact of PAD4 pharmacological inhibition on DC responses *in vitro*, our initial step was to investigate the role of mediators capable of inducing DC responses in our COVID-19 model. As previously demonstrated, and confirmed in our hands, SARS-CoV-2 does not induce the maturation of murine DCs *in vitro* (Supplementary Figure 1) [43]. Since our COVID-19 model elicits a robust TNF-α response (Figure 1H) [14,44], and given that this cytokine is an important DC stimulator in response to viral infections [45], we evaluated whether TNF-α induces DC responses in this model. We employed an antibody capable of blocking TNF-α-induced DC maturation *in vitro* (Supplementary Figure 2A-C) in our COVID-19 model (Supplementary Figure 2D). Anti-TNF-α reduced the accumulation (Supplementary Figure 2E-F) and maturation (Supplementary Figure 2G) of cDCs analyzed at 2 dpi, which resulted in a lower number of proliferating T cells in the mLNs at 6 dpi (Supplementary Figure 2H-I). These results suggest that TNF-α, at least partially, mediates cDC responses in this model.

Considering that TNF-α influences DC responses in our COVID-19 model and given the expression of PAD4 by these cells (Figure 5A-B) [46,47], we analyzed whether GSK484 could influence TNF-α-induced DC responses *in vitro*. The drug has minimally impacted TNF-α-induced maturation of BMDCs (Supplementary Figure 3A-D) and splenic DCs (Supplementary Figure 3E-F), as evidenced by MHC-II, CD80, and CD86 expression. However, we noticed that GSK484 enhanced the secretion of IL-6, IL-23, and IL-1β (Supplementary Figure 3G), while it diminished IL-10 production. Although the analysis of cytokine secretion suggests that the drug might modulate these cells to exhibit more pro-inflammatory functions, our findings reveal that GSK484 diminishes BMDC antigen presentation (Figure 5C-E), without affecting cell viability (Figure 5F). Similarly, the drug also affects antigen presentation by splenic cDC1 (Figure 5G-H), and cDC2 (Figure 5I-J). This reduction in antigen presentation appears to result from impaired antigen uptake, as GSK484 diminished the uptake of OVA by BMDCs (Figure 5K-L). Importantly, Cl-Amidine, a pan-PAD inhibitor, also diminishes antigen presentation by BMDCs (Supplementary Figure 4A-D) [46] and splenic DCs (Supplementary Figure 4E-F), suggesting that PAD4 inhibitors can affect DC functions, with potential implications for T cell priming.

**Figure 5.**
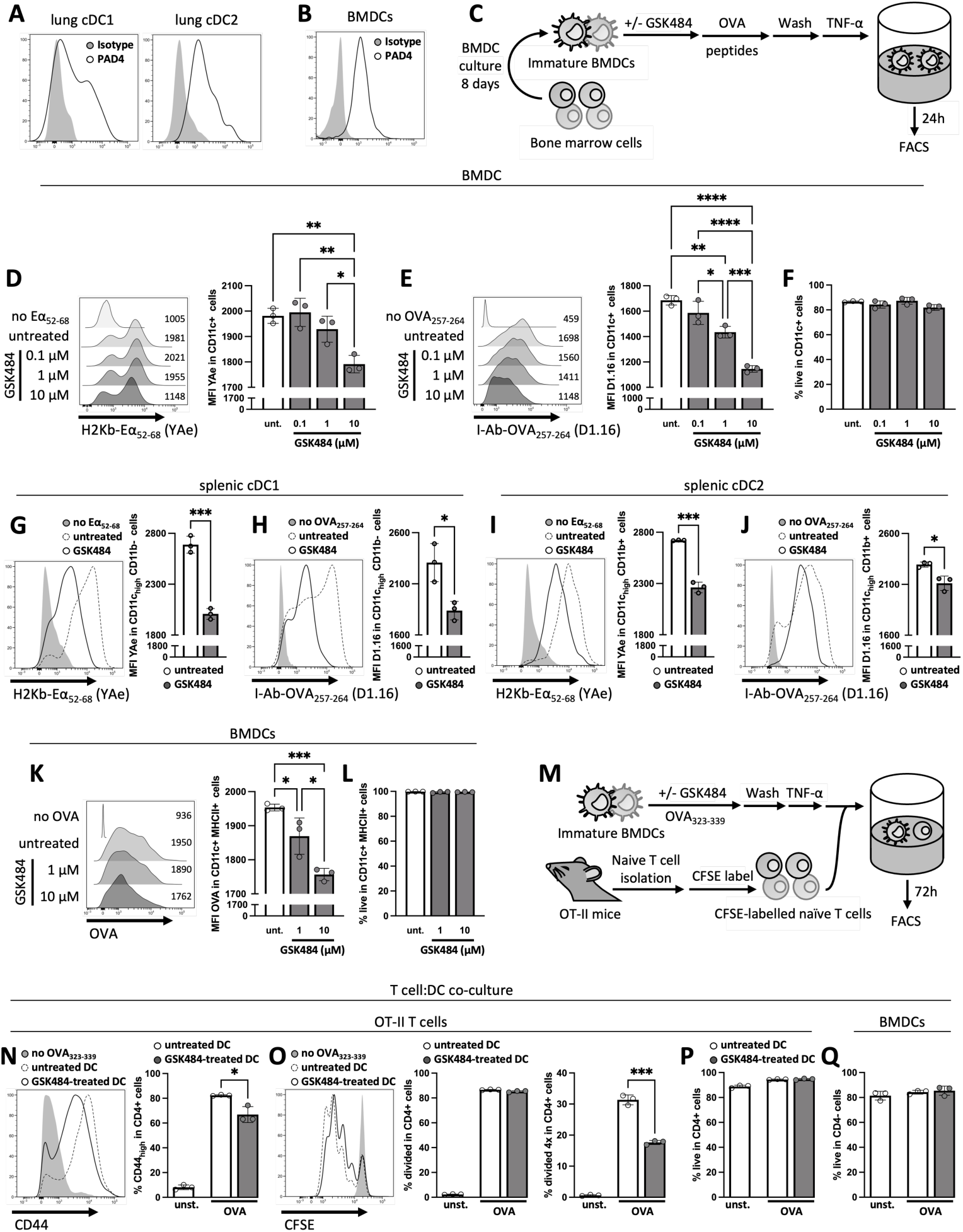
GSK484 reduces antigen presentation by impairing antigen uptake thereby hindering T cell priming. (**A**) PAD4 expression in lung CD11c_high_ MHC-II+ CD103+ CD11b-(cDC1) and CD11b+ (cDC2) cells. (**B**) PAD4 expression in BMDCs. (**C**) GSK484-treated BMDCs were pulsed with Eα_52-68_ or OVA_257-264_ for 1 hour, washed, and stimulated with TNF-α for 24 hours. (**D**) H2Kb-Eα_52-68_ (YAe) and (**E**) I-Ab-OVA_257-264_ (D1.16) median fluorescence intensity (MFI) in CD11c+ BMDCs. (**F**) Percentage of live in CD11c+ BMDCs. (**G**) YAe and (**H**) D1.16 MFI in splenic CD11c_high_ CD11b-DCs. (**I**) YAe and (**J**) D1.16 MFI in splenic CD11c_high_ CD11b+ DCs. (**K**) OVA MFI in GSK484-treated CD11c+ MHC-II+ BMDCs pulsed with OVA-AF647 for 1 hour. (**L**) Percentage of live in CD11c+ MHC-II+ BMDCs. (**M**) GSK484-treated BMDCs were pulsed with OVA_323-339_, washed, stimulated with TNF-α and co-cultured with CFSE-labelled CD4 T cells from OT-II mice. (**N**) Percentage of CD44_high_, and (**O**) cells that have divided or that have divided 4 times in CD4+ cells. (**P**) Percentage of live in CD4+ or (**Q**) CD4-cells. Differences in D-F and K-L were determined by one-way ANOVA, followed by Tukey’s multiple comparison test. Differences in G-J and N-Q were determined by unpaired t-test. *p<0.05, **p<0.01, ***p<0.001, and ****p<0.0001.

To determine whether GSK484’s effect on antigen uptake and presentation could impact T cell priming, we treated BMDCs only during the antigen uptake phase, before culturing them with CD4 T cells from OT-II mice (Figure 5M). As GSK484 is a reversible inhibitor [24], it is expected that the effects of PAD4 inhibition do not impact subsequent DC functions during the assay, such as cell maturation. GSK484-treated DCs hindered T cell priming, evidenced by a reduced percentage of CD44_high_ cells (Figure 5N) and of cells that have divided four times (Figure 5O). Importantly, the viability of T cells (Figure 5P) and BMDCs (Figure 5Q) was not affected. Taken together, these findings suggest that PAD4 targeting reduces antigen presentation by impairing antigen uptake, resulting in diminished T cell priming.

GSK484 impairs IL-2 signaling and reduces T cell proliferation

After demonstrating that PAD4 inhibitors impact DC antigen presentation, thereby leading to reduced T cell priming *in vitro*, we investigated whether this drug could directly influence T cell responses, as these cells also express PAD4 [48]. Given the reduction in the number of dividing T cells from the mLNs of mice under GSK484 treatment in our COVID-19 model (Figure 3N), we investigated the GSK484’s potential impact on T cell proliferation *in vitro*. To distinguish from an indirect effect originating from diminished antigen presentation, we employed an APC-independent system, in which T cell receptor (TCR) signalling was normalized by stimulating T cells with an equal concentration of anti-CD3. Moreover, due to the short lifespan of murine naïve T cells, discriminating between treatment-induced cell death and the potential drug’s effect on activation/proliferation would be difficult. Hence, naïve human T cells were utilized, as they also express PAD4 [49] and have a lifespan 40-50 times longer than naïve murine T cells [50]. Using a polyclonal antibody that specifically reacts with intrapeptidic citrulline, independent of the amino acid sequence, but not with free citrulline or arginine, we demonstrate that GSK484 inhibits TCR-induced protein citrullination in CD4 T cells (Figure 6A). In addition, the drug treatment did not affect T cell viability (Figure 6B) or activation (Figure 6C). However, GSK484 reduced CD4 T cell proliferation (Figure 6D). These effects were also seeing in CD8 T cells (Figure 6E-H). Our findings suggest that while GSK484 does not affect T cell activation, it markedly decreases T cell proliferation, potentially contributing to the observed reduction in virus-specific T cell responses following GSK484 treatment in our COVID-19 model.

**Figure 6.**
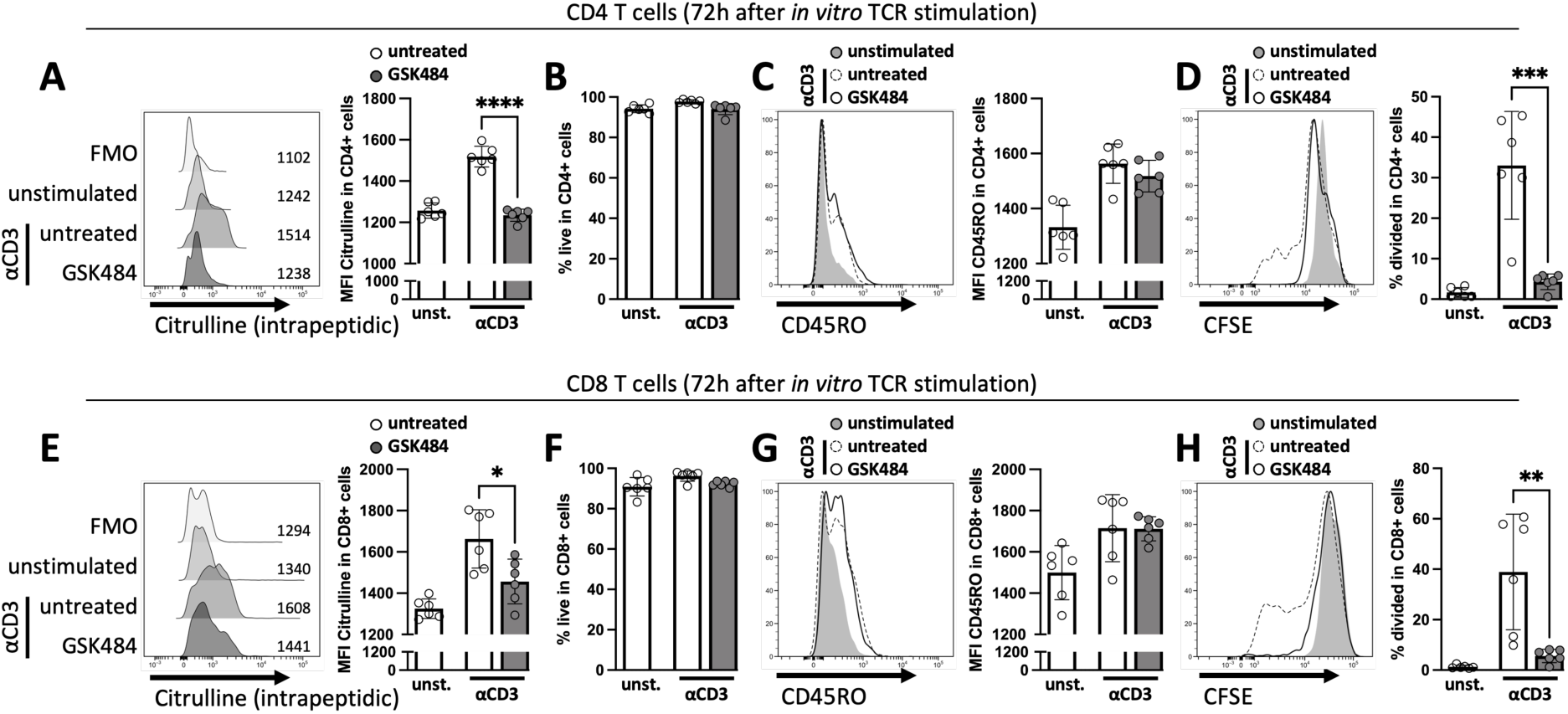
GSK484 diminishes human T cell proliferation. (**A-D**) CFSE-labelled human CD4 and (**E-H**) CD8 T cells were stimulated *in vitro* with anti-CD3 and harvested 3 days later for flow cytometry analysis. (**A**) Intrapeptidic citrulline median fluorescence intensity (MFI), (**B**) percentage of live, (**C**) CD45RO MFI, and (**D**) percentage of CFSE_dim_ in CD4+ cells. (**E**) Intrapeptidic citrulline MFI, (**F**) percentage of live, (**G**) CD45RO MFI, and (**H**) percentage of CFSE_dim_ in CD8+ cells. Differences were determined by unpaired t-test. *p<0.05, **p<0.01, ***p<0.001, and ****p<0.0001.

As GSK484 diminished T cell proliferation without affecting cell activation, we investigated whether the treatment could impact IL-2 production and signaling, as this cytokine enables T cells to survive consecutive division cycles [51]. Consistent with our previous assay, GSK484 inhibited protein citrullination (Figure 7A) without affecting activation markers such as CD44 (Figure 7B) and CD25 (Figure 7C), which is the α-chain of the IL-2 receptor. Notably, GSK484 diminished IL-2 concentration in CD4 T cell cultures 24 hours after anti-CD3/CD28 stimulation (Figure 7D), without compromising cell viability (Figure 7E). Since cell division is typically not observed in CD4 and CD8 T cells during the initial 24 hours of TCR stimulation [52], our results suggest that GSK484 affects IL-2 production, independently of cell division. This was confirmed by a decrease in the proportion of IL-2 producers in CD4 T cells (Figure 7F), resulting in impaired signaling of pSTAT5 (Figure 7G-H), which translocate to the nucleus, where it binds to target genes essential for T cell survival and proliferation [53]. Similar effects in IL-2 production and signaling were also observed in CD8 T cells treated with GSK484 (Figure 7I-P), as well as in T cells treated with Cl-Amidine (Supplementary Figure 5). These findings suggest that PAD4 inhibitors diminish T cell proliferation, at least partially, by affecting IL-2 production by these cells.

**Figure 7.**
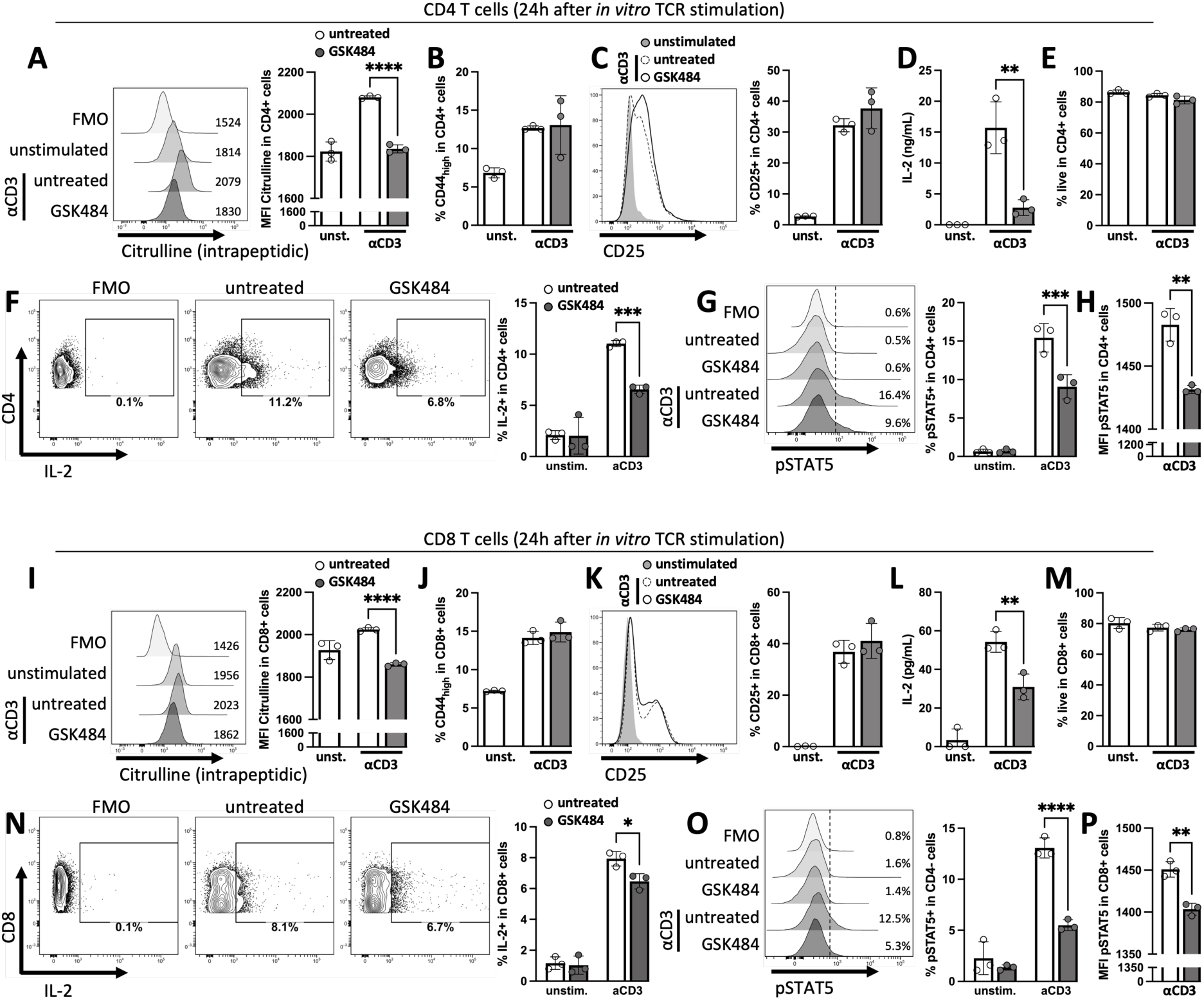
GSK484 impairs IL-2 signaling by reducing its production by T cells. (**A-H**) CFSE-labelled CD4 and (**I-P**) CD8 T cells were stimulated *in vitro* with anti-CD3 and anti-CD28 and harvested 24 hours later for analysis. (**A**) Intrapeptidic citrulline median fluorescence intensity (MFI), (**B**) percentage of CD44_high_, and (**C**) percentage of CD25+ in CD4+ cells. (**D**) Concentration of IL-2 in CD4 T cell culture supernatants. (**E**) Percentage of live, (**F**) IL-2+, and (**G**) pSTAT5+ in CD4+ cells. (**H**) pSTAT5 MFI in CD4+ cells. (**I**) Intrapeptidic citrulline MFI, (**J**) percentage of CD44_high_, and (**K**) percentage of CD25+ in CD8+ cells. (**L**) Concentration of IL-2 in CD8 T cell culture supernatants. (**M**) Percentage of live, (**N**) IL-2+, and (**O**) pSTAT5+ in CD8+ cells. (**P**) pSTAT5 MFI in CD8+ cells. Differences in A-E and H-L were determined by unpaired t-test. Differences in F-G and M-N were determined by one-way ANOVA, followed by Tukey’s multiple comparison test. *p<0.05, **p<0.01, ***p<0.001, and ****p<0.0001.

### GSK484 diminishes while rhDNase potentiates antigen-specific T cell responses in a model of lung inflammation

Considering the impact of NET targeting on DCs and T cells in the COVID-19 model and *in vitro*, we analyzed the effects of NET-targeted drugs on the activation and proliferation of adoptively transferred antigen-specific T cells in a model of acute lung inflammation (Figure 8A). In this model, we used LPS, known for its capacity to primarily stimulate DC responses via TNF-α [54,55]. GSK484 treatment inhibited protein citrullination of OT-II cells (Figure 8B), indicating a direct impact of the drug on these cells. Consistently, GSK484 reduced the number of OTII cells (Figure 8C-D), memory effector T cells (Figure 8E-F), and divided cells (Figure 8G-H) in the mLNs of animals instilled with antigen. Interestingly, rhDNase treatment enhanced these cell numbers. Previous studies have shown that DNase-fragmented, but not integrate, NETs promote DC maturation [13]. Additionally, free histones, a component of fragmented NETs (fNET), can reduce the threshold of T cell activation [11]. We show that NET incubation with rhDNase for 30 minutes is sufficient to generate fNETs that enhance T cell activation *in vitro* (Supplementary Figure 6). Altogether, our results highlight the suppressive effects of PAD4 targeting on T cell responses, while also shedding light on the potential of rhDNase treatment to boost T cell responses in tissues rich in NETs.

**Figure 8.**
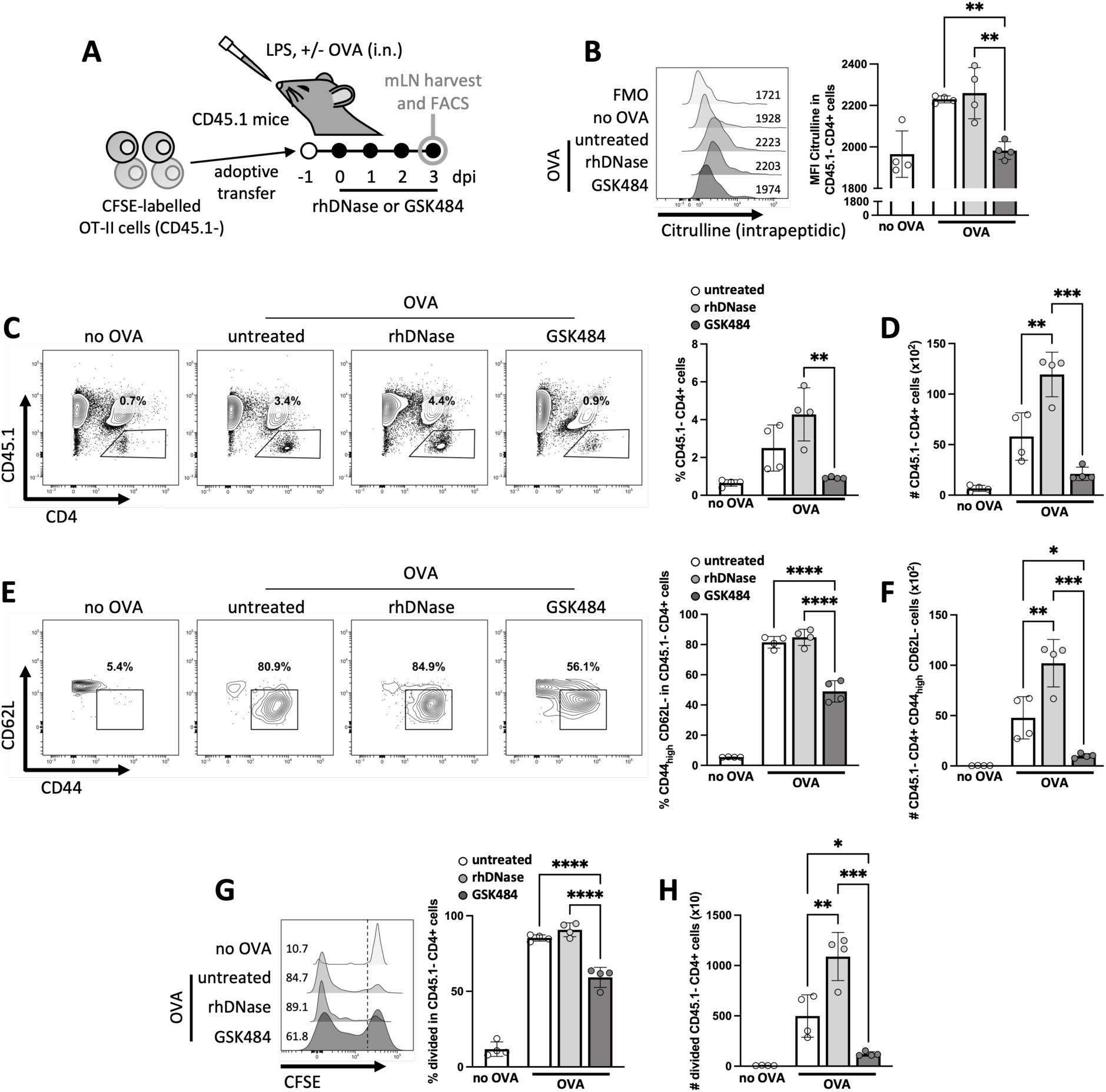
GSK484 and rhDNase exhibit contrasting effects on antigen-specific T cell activation and proliferation in a model of lung inflammation. (A) CFSE-labeled cells from lymph nodes (LNs) and spleens of OT-II mice were adoptively transferred to CD45.1 mice that were intranasally challenged with LPS and OVA and treated daily with rhDNase or GSK484. Mediastinal LNs (mLNs) were harvested 3 days later and analyzed by flow cytometry. (B) Intrapeptidic citrulline median fluorescence intensity (MFI) in CD45.1-CD4+ cells. (C) Percentage and (D) number of CD45.1-CD4+ cells. (E) Percentage of CD44_high_ CD62L-in CD45.1-CD4+ cells and (F) number of CD45.1-CD4+ CD44_high_ CD62L-cells. (G) Percentage of CFSE_dim_ in CD45.1- CD4+ cells and (H) number of CD45.1- CD4+ CFSE_dim_ cells. Differences between OVA-challenged groups were determined by one-way ANOVA, followed by Tukey’s multiple comparison test. *p<0.05, **p<0.01, ***p<0.001, and ****p<0.0001.

## Discussion

Despite the current phase of the COVID-19 pandemic being characterized by widespread immunity from prior infection or vaccination, the necessity for effective treatments persists [1]. The present study and others emphasize the importance of therapeutic strategies that not only alleviate clinical symptoms but also consider their impact on adaptive immunity [3]. NETs have been implicated in COVID-19 pathogenesis, leading to exploration of NET-targeted therapies [9,14]; however, the effects of these therapies in SARS-CoV-2-specific responses were unclear. Herein, we show that rhDNase, which degrades NET DNA, and GSK484, which inhibits PAD4-dependent NETosis, reduce lung NET concentration and improve clinical outcomes. We provide evidence that rhDNase does not affect SARS-CoV-2-specific T cell responses, key in SARS-CoV-2 clearance [4,5], while GSK484 diminish them. This reduction was attributed to GSK484’s impairment of antigen presentation by DCs and reduction of T cell proliferate response by affecting IL-2 production and signaling. In line with these observations, GSK484 diminished antigen-specific T cell responses in a model of lung inflammation. Conversely, rhDNase treatment enhanced these responses, potentially by increasing the presence of fNETs, which lower the threshold for T cell activation. These findings highlight the complex interplay between therapeutic interventions, NETs, and PAD4-expressing cells such as DCs and T cells in the context of COVID-19, providing valuable insights into potential treatment strategies.

Given the role of NETs in mediating DC and T cell responses [10–13], and considering that PAD4 is expressed by both murine and human DCs [46,47,56,57] as well as murine and human naive and memory T cells [49,58], we explored the impact of rhDNase and GSK484 on T cell immunity during SARS-CoV-2 infection. We observed that GSK484 treatment during the infection altered the responses of cDCs, which capture and present viral antigens to T cells. GSK484 limited lung cytotoxic T, Th17 and T_reg_ cell responses, while also reducing the accumulation of dividing T cells in lung draining LNs. Notably, GSK484, but not rhDNase, particularly inhibited SARS-CoV-2-specific T cell effector responses. This inhibitory effect aligns with other studies where PAD4 inhibition, through Cl-Amidine or genetic depletion, reduced Th1, T_reg_, and/or Th17 activity in various disease models, such as non-alcoholic steatohepatitis [59], DSS-induced colitis [60], and type 1 diabetes [61]. Our findings underscore the contrasting effects of rhDNase and GSK484 on T cell responses during SARS-CoV-2 infection.

In our *in vitro* assays to assess the effects of PAD4 inhibitors on DCs, we observed that GSK484 minimally impacts DC maturation induced by TNF-α, a cytokine we demonstrated to mediate DC responses during SARS-CoV-2 infection. However, the drug enhanced pro-inflammatory cytokine secretion. Although these results suggest GSK484’s potential to enhance DC pro-inflammatory functions, our results demonstrate that GSK484 impairs antigen uptake and diminishes MHC-I and MHC-II-mediated antigen presentation by these cells. Consistently, we show that this impaired antigen presentation leads to reduced antigen-specific T cell priming. Moreover, we and others have shown that Cl-Amidine, another PAD inhibitor, disrupts DC antigen presentation, resulting in impaired CD4 and CD8 T cell proliferation *in vitro* [46]. In contrast, a recent report utilizing cells from myeloid cell type-specific *Padi4* conditional knockout mice indicates increased MHC-II expression on macrophages from these animals, resulting in enhanced expression of antigen presentation gene signatures [48]. Confirming this trend within the same mouse lineage, we found that BMDCs from mice lacking PAD4 also exhibit higher levels of MHC-II compared to their controls, resulting in enhanced antigen presentation *in vitro* (unpublished data). The divergent outcomes in antigen presentation between cells treated with GSK484 or Cl-Amidine and those from animals lacking PAD4 may be attributed to off-target effects of the drugs. Alternatively, the high level of PAD4 expression in mouse bone marrow cells, its role in hematopoiesis [62], and its involvement in the differentiation of other cell types, such as cardiomyocytes and endothelial cells [63], might suggest a role for PAD4 in the differentiation of APCs. This hypothetical impact could potentially drive the differentiation of APCs lacking PAD4 towards subsets with increased MHC expression and enhanced capacity for antigen presentation [64–66]. While indicating the need for further investigation into the precise mechanisms by which GSK484 and Cl-Amidine affect DC responses *in vitro*, our findings underscore their impact on antigen uptake and presentation, ultimately affecting T cell responses.

While we demonstrate that impaired T cell priming can result from GSK484’s impact on DC antigen presentation, we also provide evidence that this drug can directly influence T cell responses. Under *in vitro* TCR signaling, we observed a significant reduction in the proliferation of human CD4 and CD8 T cells with GSK484 treatment, while activation markers remained unaffected. Previous studies have reported that PAD4-selective inhibition with GSK199 does not affect anti-CD3/CD28-mediated activation of human CD4 T cells [49]. Additionally, another study indicates that both CD4 and CD8 T cells from mice lacking PAD4 exhibit significantly reduced proliferative capacity compared with cells from wild-type mice [67]. As an inhibitor of PAD4, an enzyme catalyzing protein citrullination, we found that GSK484 diminishes intracellular protein citrullination in mouse and human CD4 and CD8 T cells. This aligns with literature reports of diminished citrullination of histone H3 in mouse [68] and human T cells [69] under treatment with Cl-Amidine, as well as in PMA-stimulated T cells from mice lacking PAD4 [69]. Citrullination can alter the function of diverse proteins, such as NF-κB p65, promoting its nuclear translocation [70], a crucial step for T cell proliferation and effector/memory functions [71,72]. One of the functions regulated by NF-κB activation is IL-2 production [73]. During the first phase of the cell cycle that allows cells to proliferate, activated T cells require survival/stimulatory signals, such as IL-2, to enter the expansion phase and divide rapidly [51,74]. We found that both GSK484 and Cl-Amidine decreased the secretion of IL-2 by activated T cells. IL-2 signals through JAK1/3 to phosphorylate STAT5 and ultimately drive T cell proliferation [75]. Accordingly, the levels of pSTAT5 were also diminished by treatment with the PAD4 inhibitors. It is worth noting, however, that GSK484 and Cl-Amidine might affect T cells differently, as the latter drug also inhibits other PAD enzymes, such as PAD2, which controls Th2/Th17 balance in murine [49,68] and human [69,76] cells. Overall, these results suggest that pharmacological inhibition of PAD4 expressed by T cells suppresses T cell proliferation, at least partially by reducing IL-2 production and, consequently, IL-2 signaling.

Analyzing the impact of rhDNase and GSK484 on adoptively transferred antigen-specific T cell responses in a lung inflammation model revealed that GSK484 diminished T cell activation and, while rhDNase potentiated these effects. Although both DNase treatment and PAD4 inhibition can prevent tissue damage, it has been demonstrated that, unlike PAD4 inhibition, DNase might not effectively prevent the accumulation of NET-derived components in inflamed tissues, such as histones [77]. This suggests that DNase might increase the availability of free histones by degrading NET backbone. Histones account for approximately 70% of all NET-associated proteins [78], and free histones were found to enhance T cell activation *in vitro* and *in vivo* [11]. In addition, DNase-fragmented NETs, but not integrate NETs were found to induce murine and human DC maturation [13]. This stimulatory effect is also believed to be induced by NET histones, as free histones were found to stimulate pro-inflammatory cytokine production by BMDCs [79]. In this present work, we show that DNase-fragmented NETs, but not intact NETs, directly potentiate T cell activation. In line with these results, a report showed that DNase treatment can prevent the reduction of T cell accumulation in lymphoid organs of ischemically injured mice [80]. Furthermore, DNase treatment or intratracheal administration of DNase-fragmented NETs were found to induce the activation of naive CD4 T cells adoptively transferred into lung allograft recipients, as well as to stimulate intragraft effector T cell accumulation [13]. Differences in endogenous DNase secretion might account for the variations observed between the COVID-19 model and the lung acute inflammation model shown in this present work. Additionally, a potential difference in the concentration of cytokines such as IL-6, which was reduced in the COVID-19 model and plays a role in T cell responses [81], could also contribute to these variations. Nevertheless, these results suggest that while PAD4 inhibitors can diminish antigen-specific T cell responses *in vivo*, rhDNase may preserve or enhance these responses in tissues rich in NETs by modulating the availability of NET components in the microenvironment.

In summary, this study elucidates the complex interplay between NETs and PAD4 in shaping adaptive immune responses during SARS-CoV-2 infection. By comparing rhDNase and GSK484 treatments, we demonstrate how NET degradation and PAD4 targeting can differentially shape immune outcomes. The ability of rhDNase to potentiate T cell responses *in vitro* and *in vivo* suggests its potential utility in therapeutic strategies aimed at maintaining or enhancing virus-specific immunity while diminishing the deleterious effects of NETs. Meanwhile, the suppressive effects of PAD4 inhibitors call for a careful evaluation of their use, particularly in contexts where robust T cell immunity is crucial. While adaptive immune responses in COVID-19 are relevant for combating the virus, they might also be responsible for over-inflammation and pathology [82,83]. As such, similar to steroids, which were widely used during the pandemic to control over-inflammation but resulted in weaker SARS-CoV-2-specific T cell responses [84], PAD4 inhibitors might still be useful for controlling over-inflammation. In this context, studies on the influence of NET-targeted drugs in models of long COVID-19 are essential to evaluate their impact in immunopathology caused by potential altered T cell responses. Future research could also explore potential combinatory treatments that can modulate adaptive immunity while preserving or increasing antiviral efficacy. By enhancing our understanding of the interplay between NETs, therapeutic interventions, and adaptive immunity this study provides a foundation for developing more effective and balanced treatment strategies for COVID-19 and other diseases involving excessive NET formation.

## Author contributions

CSB, FPV, and FQC designed the COVID-19 model experiments. CSB, FPV, GFG, and RMRL conducted the COVID-19 model experiments. CSB designed and conducted the remaining experiments, acquired data, and analyzed data. ASR, EA, JACF, TMC, and FQC provided materials. FQC acquired funding and supervised the study. CSB wrote the manuscript. All authors reviewed the manuscript and provided final approval for submission.

## Acknowledgments

This work was supported by the São Paulo Research Foundation (FAPESP), grant n° 2013/08216-2, 2020/05601-6, and 2021/09539-6.

