## Supplementary material for "Differential action modes of Neutrophil Extracellular Trap-targeted drugs define T cell responses in SARS-CoV-2 infection": Suplemmentary Figures

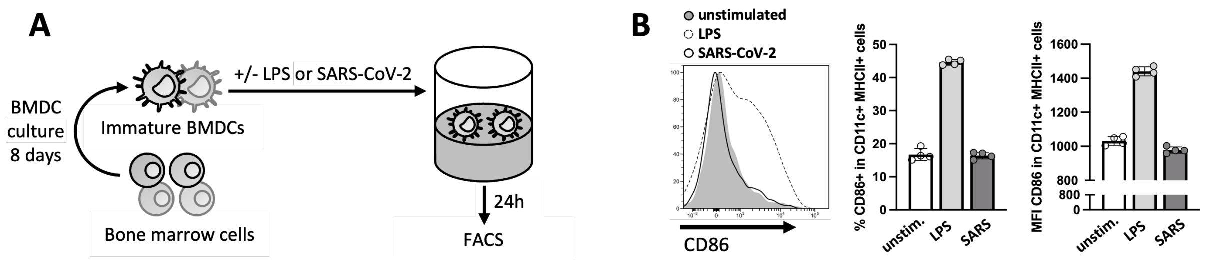


**Supplementary Figure 1**. SARS-CoV-2 does not induce murine DC maturation *in vitro*. (**A**) Bone marrow-derived dendritic cells (BMDCs) were stimulated with LPS or SARS-CoV-2 (1 PFU/cell) for 24 hours and analyzed by flow cytometry. (B) Percentage of CD86+ and CD86 median fluorescence intensity (MFI) in CD11c+ MHC-II+ cells.

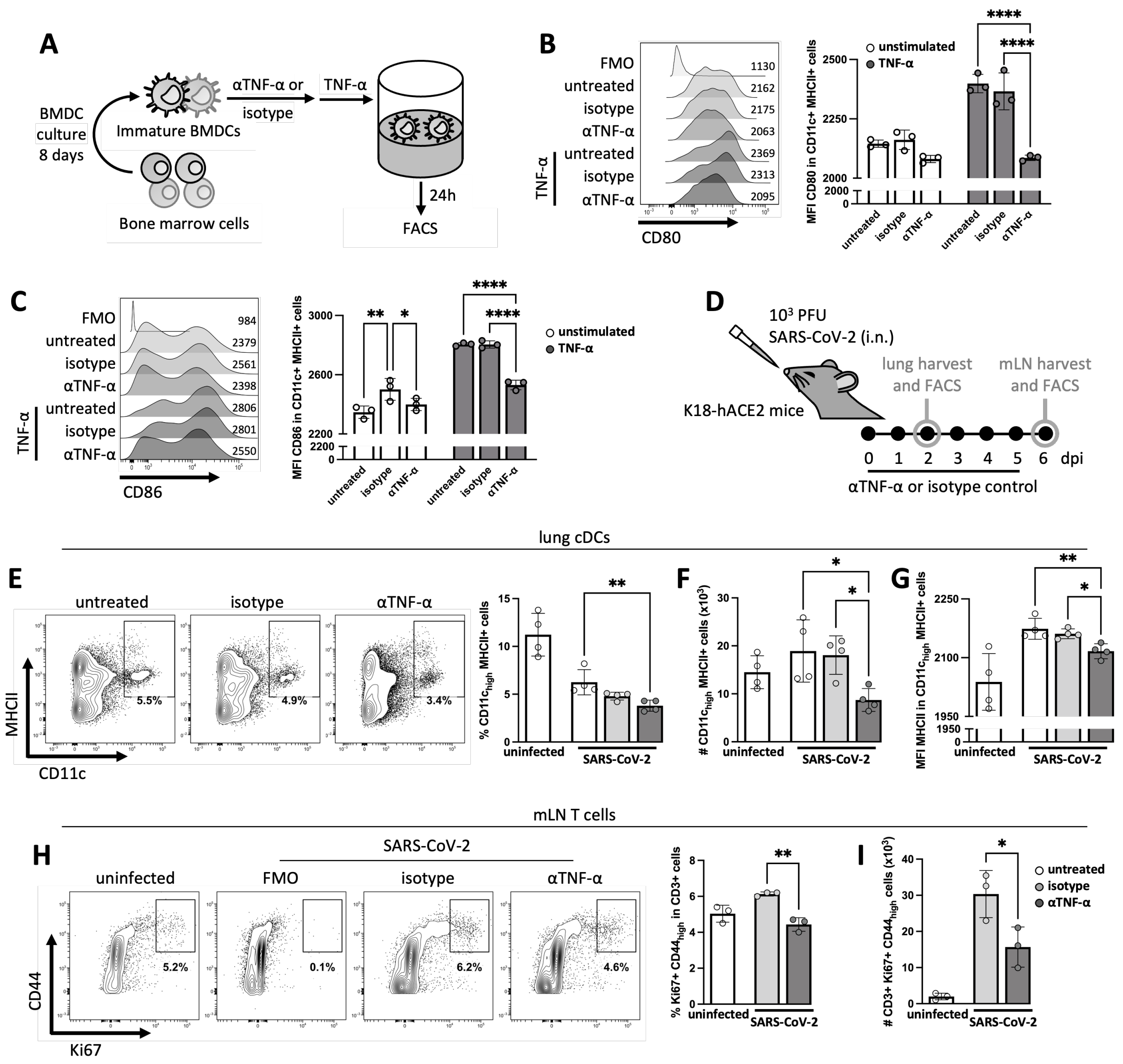


**Supplementary Figure 2**. TNF-α mediates cDC responses in the COVID-19 model. (**A**) Anti-TNF-α- or isotype control-treated BMDCs were stimulated with TNF-α for 24 hours. (**B**) CD80 and (**C**) CD86 median fluorescence intensity (MFI) in CD11c+ MHC-II+ BMDCs. (**D**) Lungs and mediastinal lymph nodes (mLN) from SARS-CoV-2-infected mice (N=3-4) treated daily with anti-TNF-α or its isotype control were harvested 2 and 6 days post-infection (dpi), respectively. (**E**) Percentage and (**F**) number of $CD11c_{high}$ MHC-II+ cells in the lungs. (**G**) MHC-II MFI in lung $CD11c_{high}$ MHC-II+ cells. (**H**) Percentage of Ki67+ $CD44_{high}$ in CD3+ cells and (**I**) number of CD3+ Ki67+ $CD44_{high}$ cells in the mLNs. Differences in B-C and E-G were determined by two-way and one-way ANOVA, respectively, followed by Tukey's multiple comparison test. Differences in H-I were determined by unpaired t-test. *p<0.05, **p<0.01, and ****p<0.0001.

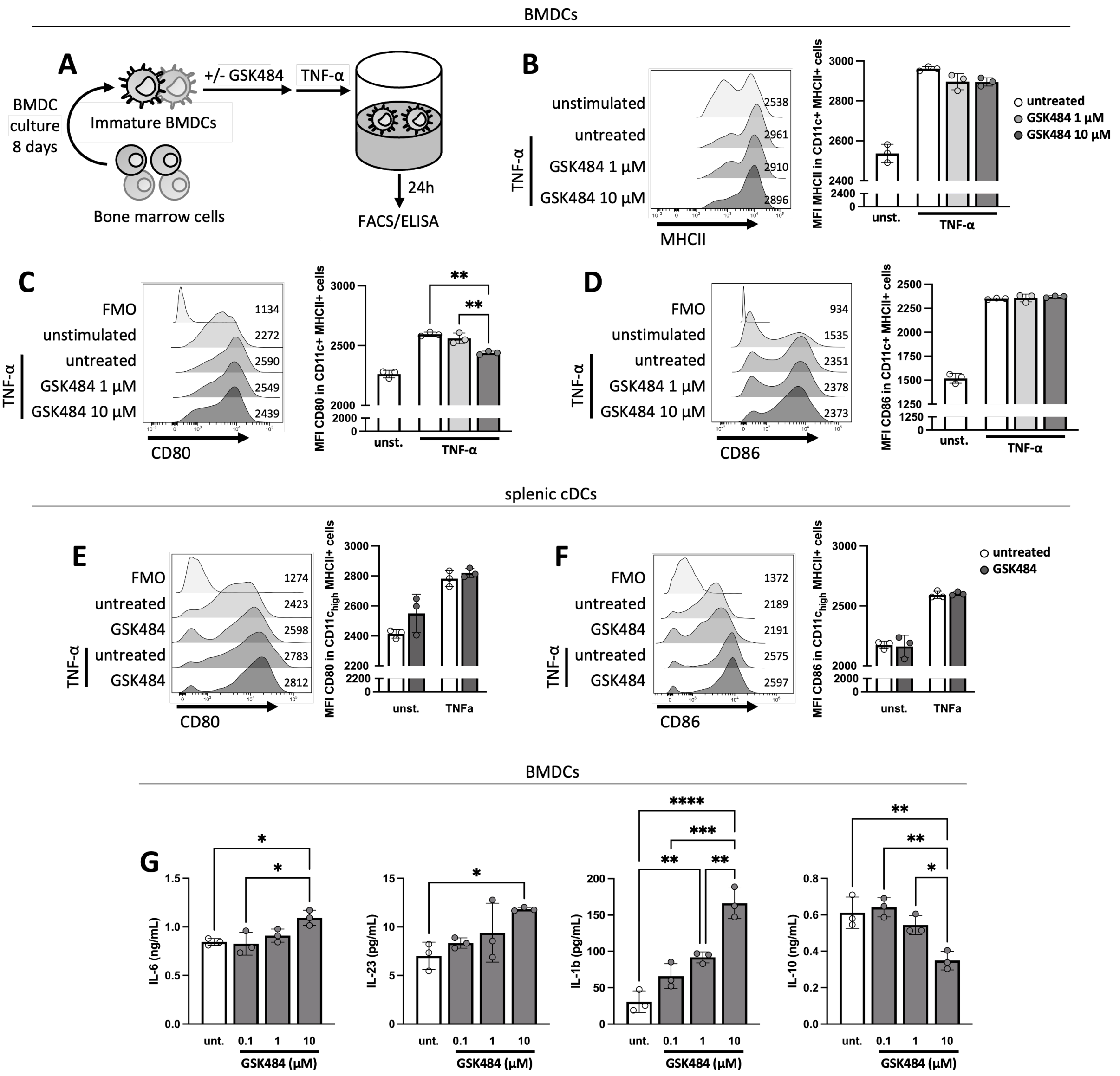


**Supplementary Figure 3.** GSK484 minimally impacts TNF-α-induced DC maturation while it alters cytokine secretion. (**A**) Bone marrow-derived dendritic cells (BMDCs) were treated with GSK484, stimulated with TNF-α for 24 hours. (**B**) MHCII, (**C**) CD80, and (**D**) CD86 median fluorescence intensity (MFI) in CD11c+ MHC-II+ BMDCs. (**E**) CD80 and (**F**) CD86 MFI in $CD11c_{high}$ MHC-II+ splenic DCs. (**G**) Concentration of cytokines from BMDC culture supernatants. Differences in B-D and G were determined by one-way ANOVA. Differences in E-F were determined by two-way ANOVA. ANOVA tests were followed by Tukey's multiple comparison test. *p<0.05, **p<0.01, ***p<0.001, and ****p<0.0001.

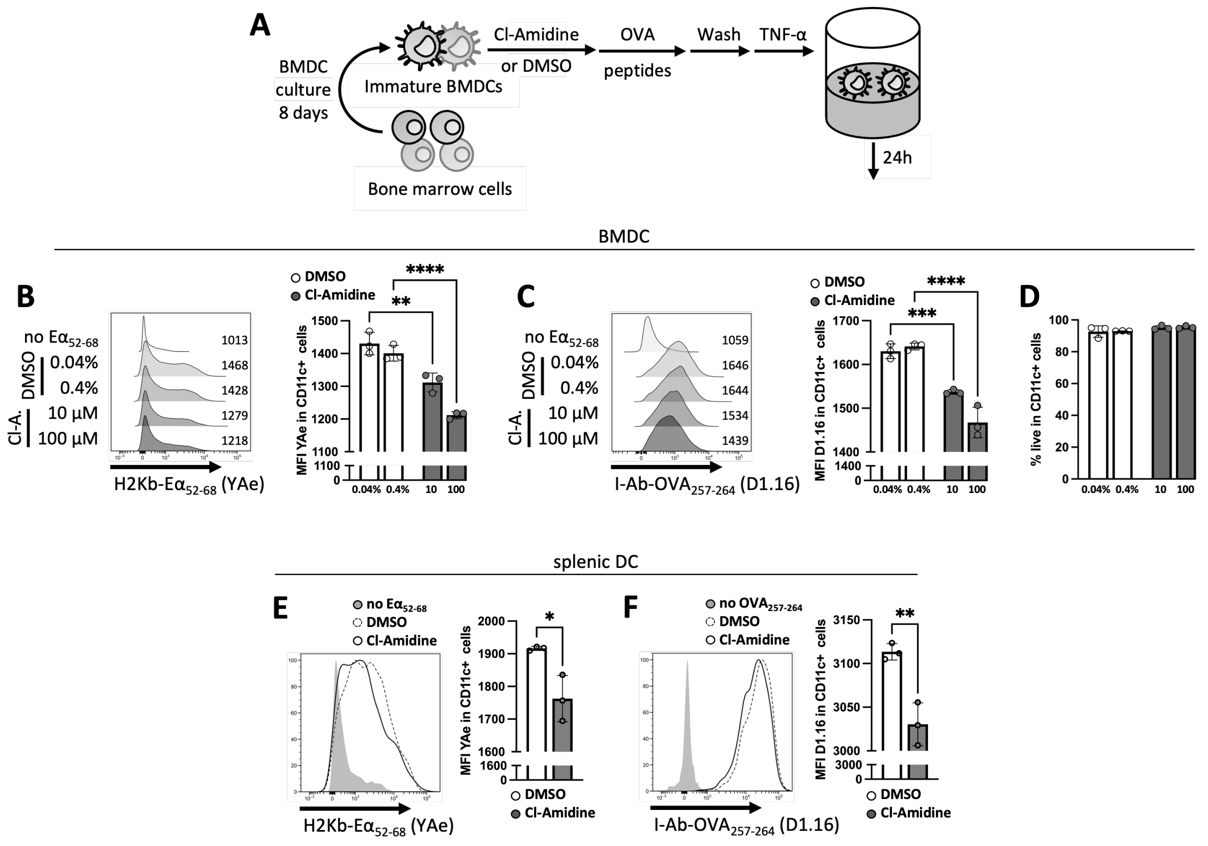


**Supplementary Figure 4**. Cl-Amidine exerts similar effects to GSK484 on DC antigen presentation. (**A**) BMDCs were treated with Cl-Amidine or equivalent concentrations of vehicle (DMSO), pulsed with $E\alpha_{52-68}$ or $OVA_{257-264}$ for 1 hour, washed, and stimulated with TNF-α for 24 hours before being analyzed. (**B**) H2Kb-$E\alpha_{52-68}$ (YAe) and (**C**) I-Ab-$OVA_{257-264}$ (D1.16) median fluorescence intensity (MFI) in CD11c+ BMDCs. (**D**) Percentage of live in CD11c+ BMDCs. (**E**) YAe and (**F**) D1.16 MFI in splenic CD11c+ DCs. Differences in B-D were determined by two-way ANOVA, followed by Tukey's multiple comparison test. Differences in E-F were determined by unpaired t-test. *p<0.05, **p<0.01, ***p<0.001, and ****p<0.0001.

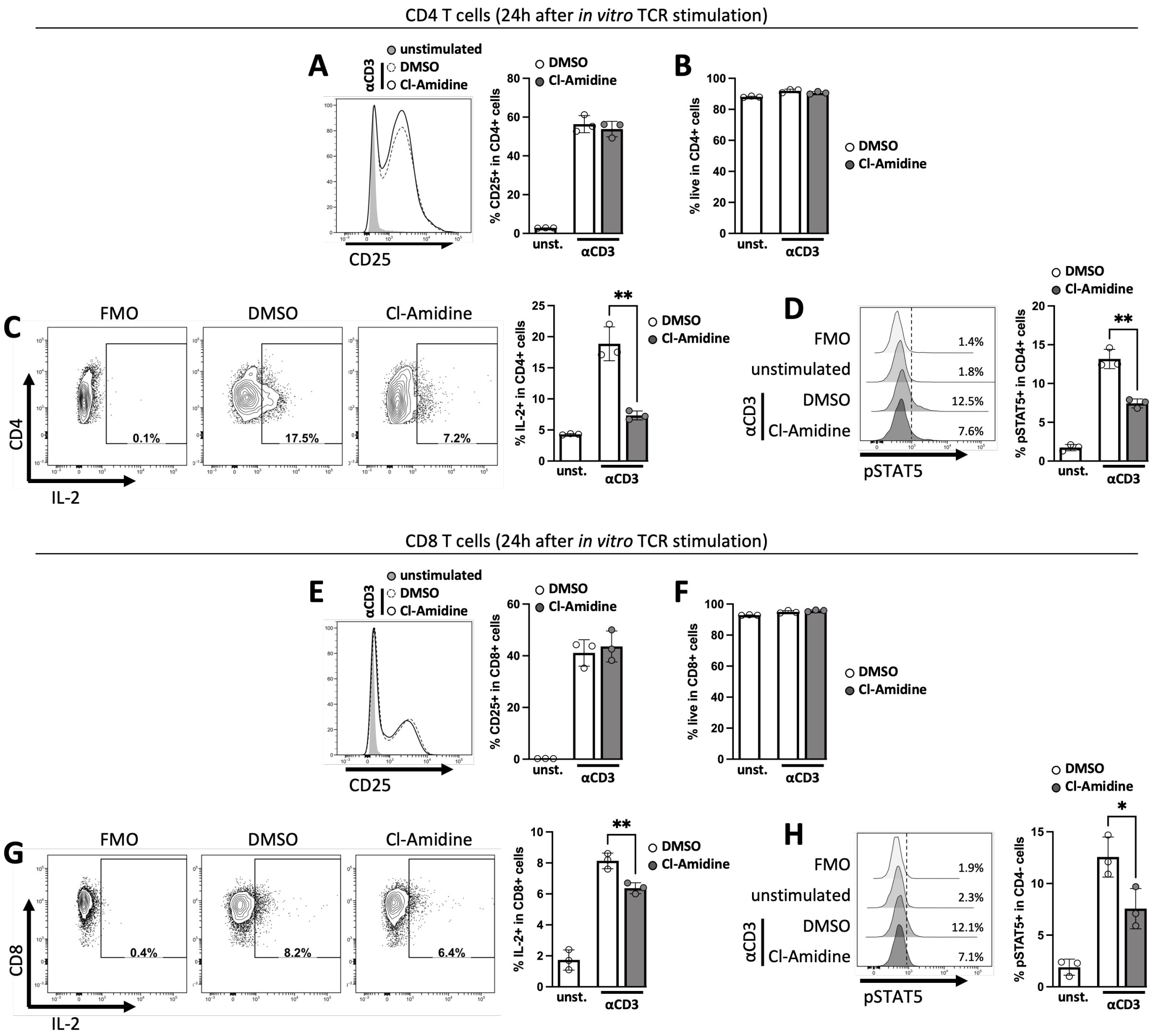


**Supplementary Figure 5**. Cl-Amidine exerts similar effects to GSK484 on T cell IL-2 signaling and production. (**A-D**) CFSE-labelled CD4 and (**E-H**) CD8 T cells were stimulated *in vitro* with anti-CD3 and anti-CD28 and harvested 24 hours later for flow cytometry analysis. (**A**) Percentage of CD25+, (**B**) live, (**C**) IL-2+, and (**D**) pSTAT5+ in CD4+ cells. (**E**) Percentage of CD25+, (**F**) live, (**G**) IL-2+, and (**H**) pSTAT5+ in CD8+ cells. Differences between stimulated groups were determined by unpaired t-test. *p<0.05, **p<0.01.

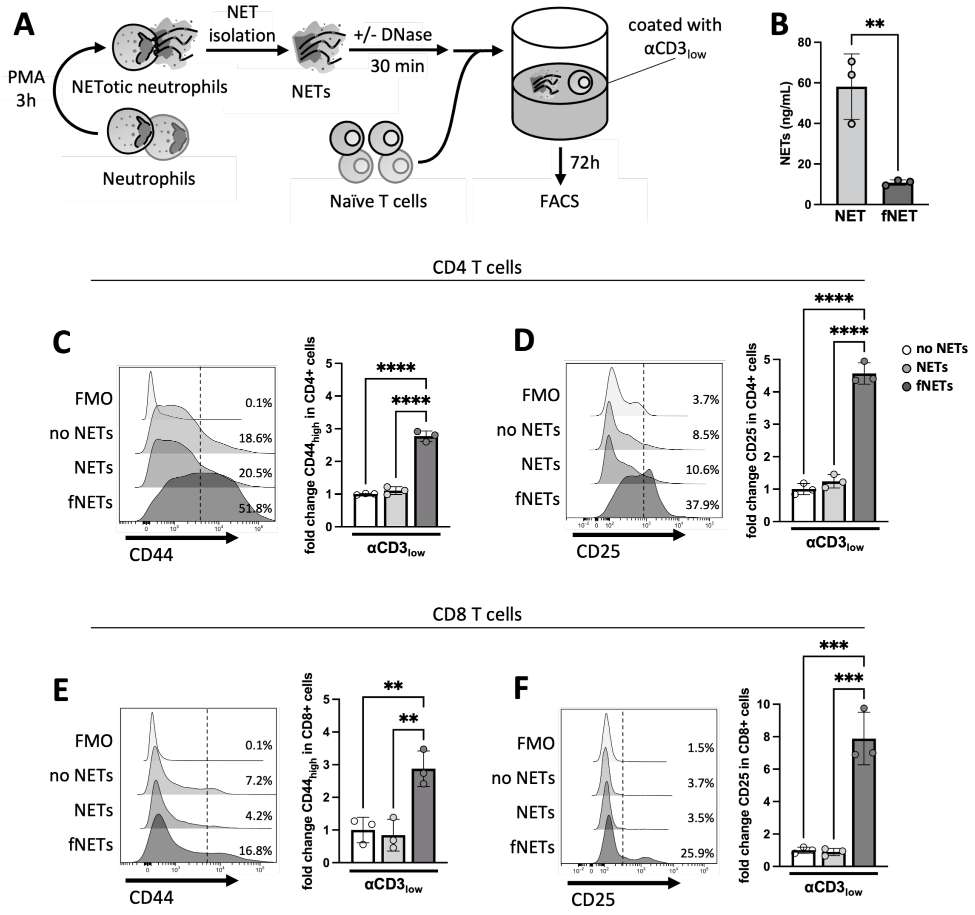


**Supplementary Figure 6**. DNase-fragmented, but not integrate NETs exert positive effects on T cell activation. (**A**) Naïve T cells were stimulated with a low concentration of anti-CD3 and co-stimulated with NETs, which were pre-treated with rhDNase or left untreated. Cells were harvested 72 hours later for flow cytometry analysis. (**B**) Concentration of NETs (MPO-DNA conjugates) in untreated (NET) or rhDNase-fragmented (fNET) NETs. (**C**) Fold change in the percentage of $CD44_{high}$ and (**D**) CD25+ in CD4+ cells. (**E**) Fold change in the percentage of $CD44_{high}$ and (**F**) CD25+ in CD8+ cells. Differences in B were determined by unpaired t-test. Differences in C-F were determined by one-way ANOVA, followed by Tukey's multiple comparison test. $^{**}p<0.01$, $^{***}p<0.001$, and $^{****}p<0.0001$.
